# Cellular inhibitor of apoptosis protein (cIAP) is a convergent hub that couples BCR–CD40 signal dynamics to B cell fate

**DOI:** 10.64898/2026.08.26.747440

**Authors:** Kentaro Inoue, Hisaaki Shinohara

## Abstract

BCR and CD40 signals are integrated to shape B cell transcriptional output, but how the two pathways are coordinated at the level of individual signaling nodes is incompletely understood. Using primary mouse B cells stimulated with anti-IgM, anti-CD40, or both (Both), we defined six DEG classes reflecting costimulation-dependence and cross-pathway antagonism. We also connected a CD40-driven, BCR-suppressed B-cell-identity module (Pax5, Aicda, Bcl6, Cd79a/b, Cd19) to an increase in Blimp1 protein at 24 h. Building on our previous work, which showed that cIAP prolongs IKK/ERK activity after single-pathway stimulation (Shinohara et al., 2016), we added an IAP inhibitor 10 min after Both stimulation and generated transcriptomic data. This profiling showed that cIAP is required not only for late canonical NF-κB-driven genes, as predicted, but also, unexpectedly, for the entire BCR-dominant, CD40-independent negative-feedback module (Cd5, Il10, Nfkbid, Spry1/2) identified in the first dataset. A separate module was instead amplified by IAP inhibition, with kinetics compatible with non-canonical NF-κB de-repression, directly addressing a question we left open in that study regarding the BCR-side partners of cIAP. Together, these correlative findings converge on a single model: cIAP is not a CD40-restricted adaptor but a hub shared by BCR and CD40, whose feedback simultaneously sustains canonical, fate-instructive signaling and restrains the non-canonical pathway, thereby coupling receptor-proximal signal dynamics to the B cell’s downstream transcriptional and fate decisions.

## Main Text

BCR engagement by antigen and CD40 engagement by CD40L on cognate T follicular helper cells are the two principal signals driving B cell activation and germinal center (GC) entry. Their relative strength and duration shape cell-fate decisions between GC recycling and plasma cell exit (*1–4*). The duration, not merely the presence, of kinase activity is itself instructive for downstream gene-expression specificity (*5, 6*). How BCR and CD40 inputs are integrated at the level of individual signaling nodes, rather than at the level of downstream transcription factors or population-level dynamics alone (*7*), remains incompletely defined. We previously showed, using single-pathway (BCR-or CD40-only) stimulation of DT40 and primary B cells, that the E3 ubiquitin ligase cIAP1 (IAP) is an early-response gene that acts as positive transcriptional feedback, prolonging IKK and ERK activity. IAP had already been implicated as a global regulator of NF-κB and MAPK activation downstream of TNF-family receptors (*8*) and as essential for normal B cell survival signaling and the GC response (*9*). Pharmacological IAP inhibition added 10 min post-stimulation, the design used throughout this study, shortened kinase activity and reduced late-phase gene expression. Genetic IAP deletion in DT40 cells confirmed an analogous requirement downstream of BCR alone. That study left three questions open: whether IAP plays the same role under physiological BCR plus CD40 costimulation (Both); whether its short-timescale (≤120 min) effects connect to longer-timescale fate outcomes; and which molecules mediate IAP’s action downstream of BCR specifically. We had explicitly flagged the last of these as unresolved (*10*). We address all three here. Underlying all three questions is a single working hypothesis: IAP may act not merely as a CD40-induced feedback regulator but as a receptor-proximal hub shared by BCR and CD40, such that its feedback dynamics, rather than either pathway’s transcription factors alone, dictate the duration and identity of the resulting gene-expression program.

We first confirmed, in our own primary mouse B cell system, that IAP inhibition added 10 min after BCR and CD40 costimulation (Both) sharply attenuates both IKK and ERK activation (Fig. 1). This result reproduces our core kinetic finding (*10*) under costimulation rather than single-pathway stimulation, and it validates the experimental paradigm used for the transcriptomic experiments below. When cells stimulated by both were compared with cells stimulated by BCR or CD40 alone (Fig. 2A), six DEG groups were identified (Figures 2B and 2C). Group B comprised costimulation-dependent induction of Myc and Tnfaip3/A20. Tnfaip3/A20 is a ubiquitin-editing enzyme that limits B-cell survival and prevents autoimmunity (*11, 12*); loss of its function exclusively in B cells leads to excessive B-cell activation and autoimmunity (*13, 14*). In the Both-vs-CD40 DEG set (focusing primarily on Group A, which accounts for the majority of the up-regulated group), in particular, the addition of anti-IgM after 30 minutes resulted in an immediate and marked enrichment of the ‘B-cell proliferation and activation pathway’, the ‘ERK1/2 cascade’ and the ‘response to mechanical stimulus’ (Fig. 2D). This is consistent with the known crosstalk mechanism whereby the BCR pathway rapidly induces additional ERK activation that cannot be achieved by CD40 alone (*15, 16*). This Group A comprised BCR-dominant induction of Cd5, Ctla4, Il10, Pdcd1lg2, Irf4, Nfkbid, and Spry1/2 (Fig. 2D,E). These findings are consistent with a stepwise induction of IRF4 dependent on the intensity of antigen receptor signaling (*17*). In Group C, which accounts for the upregulated group in the ‘Both-vs-IgM’ DEG set in Fig. 2D, the emphasis on ‘mononuclear/leukocyte proliferation’, ‘regulation of interleukin-6 production’ and ‘type II interferon production’ suggests that the addition of CD40 triggered a CD40-specific NF-κB/AP-1/C/EBP-dependent IL-6 transcription program (*18*) that cannot be elicited by BCR stimulation alone. This is a classic costimulation effect, whereby the weak proliferation and cytokine production signals induced by BCR stimulation alone are markedly enhanced by CD40 costimulation. This Group C consisted of CD40-dominant induction of Fas, Bcl2l1 and Gadd45b. It has been reported that BCR stimulation alone can induce caspase-8-dependent apoptotic signaling, and that high-dose CD40 stimulation saturates NF-κB signaling, whereby the synergistic effect of BCR can instead turn into antagonism (*7*). The appearance of ‘leukocyte apoptotic process’ and ‘regulation of DNA-binding transcription factor activity’ at the 180-minute time point in Group D (the majority of the downregulated group in the ‘Both-vs-CD40’ DEG set in Fig. 2D) suggests that the addition of BCR may have attenuated part of the survival and transcription factor program induced by CD40 alone, which is consistent with this mechanism of antagonism. Group D was a CD40-inducible, BCR-inhibitory B-cell identity module composed of core GC/B-cell identity and memory fate regulators (Pax5, Aicda, Bcl6, Cd79a/b, Cd19, Bank1, Ms4a1 (*19, 20*)) (Fig. 2D,E), and it was the largest and statistically most robust class. Two smaller inhibitory groups completed the set. The fact that Myc requires both signals is consistent with the BCR-CD40 co-dependence reported in naive B cells (*21*). The CD40-induced/BCR-inhibited identity module anticipated a marked, Both-specific increase in Blimp1 protein levels after 24 hours. Indeed, the addition of an IAP inhibitor clearly suppressed the Blimp1 protein induced by costimulation (Fig. 2F). This is broadly consistent with the model of fates, namely, “GC recycling” and “differentiation into plasma cells”,that depend on the strength of interaction with Tfh cells (*3*), as well as with the role of ERK signaling as a switch that integrates BCR-and cytokine-derived inputs to regulate plasma cell differentiation (*22*). However, bulk immunoblots cannot distinguish between changes in the homogeneous population and changes in individual subpopulations.

**Fig. 1.**
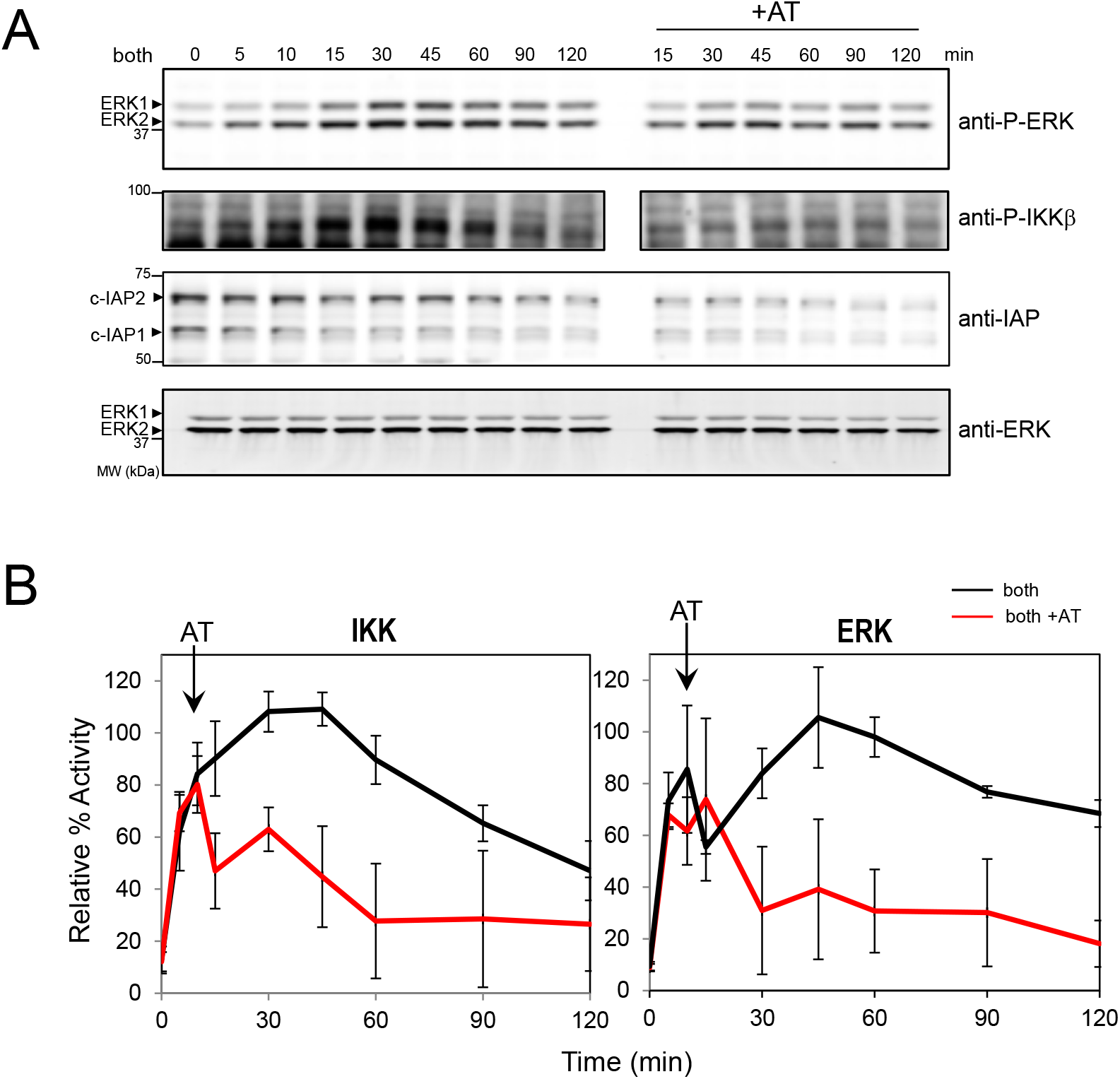
IAP inhibition attenuates IKK and ERK phosphorylation induced by BCR with CD40 costimulation. **A.** Splenic B cells (Splenic B) from mice were stimulated with 10 μg/ml anti-mouse IgM and 4 μg/ml anti-mouse CD40 (both) over the indicated time period. The IAP inhibitor added 10 min after costimulation (+AT). Whole cell lysates were analyzed by immuno blotting with anti-phospho-IKK or anti-phospho-ERK polyclonal Abs (n = 2). **B.** Quantification of IKK and ERK activity. The graph represented the quantitated phosphorylated IKK and ERK stimulated with anti-IgM and anti-CD40 (both: black line) or the IAP inhibitor added 10 min after costimulation (both+AT: red line). Data represent the means ± s.d. (n = 2).

**Fig. 2.**
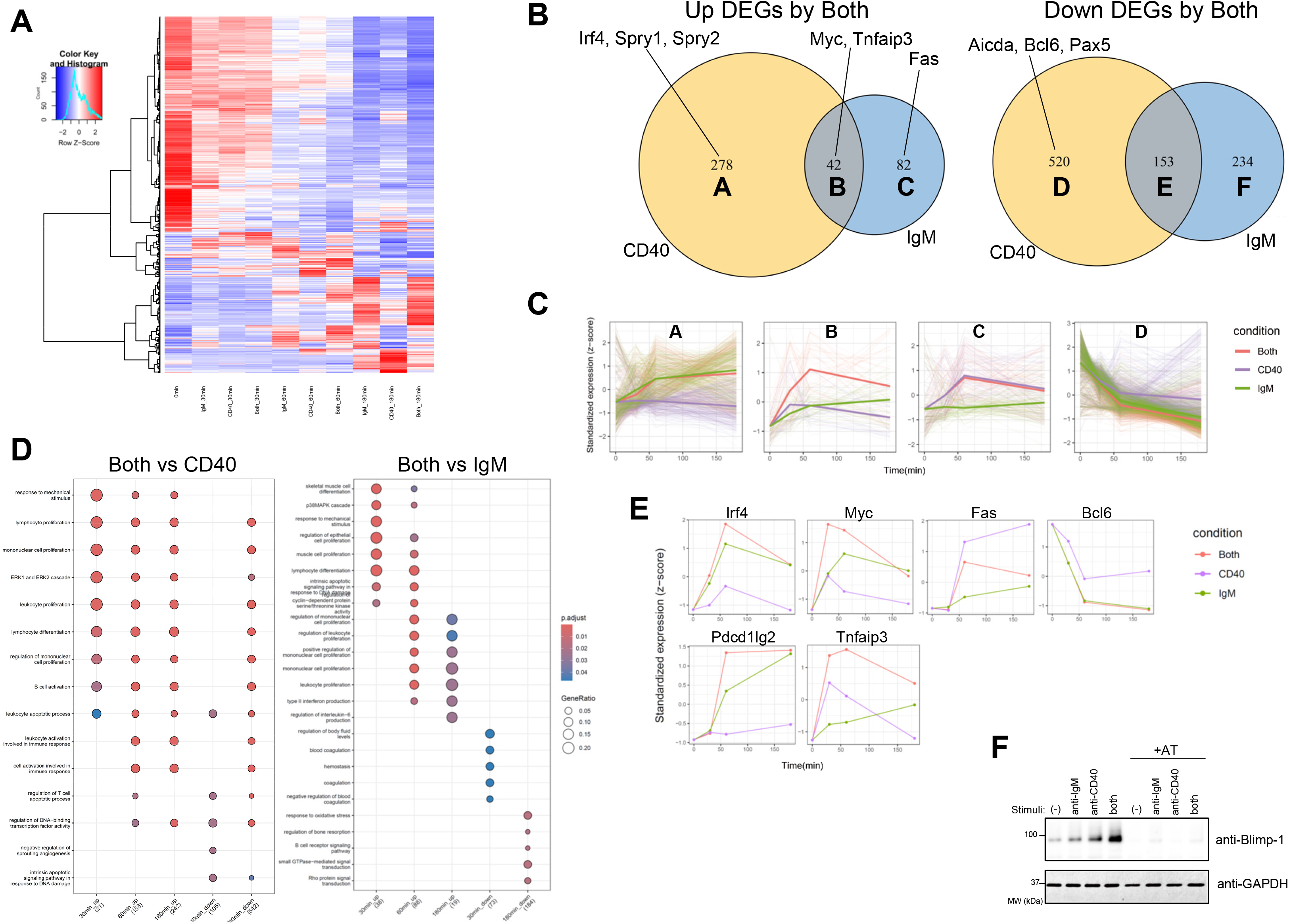
BCR and CD40 signals define six antagonistic and cooperative DEG classes that anticipate a Both-specific plasma-cell-associated transcriptional shift. **A.** Heatmap (row z-scores, n = 2) of DEGs between anti-IgM (IgM), anti-CD40 (CD40), and anti-IgM and anti-CD40 (both) treated B cells (|log2FC| > 1.0). **B.** The Venn diagram on the left (Up DEGs by Both) shows genes upregulated in costimulated (Both) cells relative to anti-CD40 alone (Group A), anti-IgM alone (Group C), or both comparisons (Group B). The Venn diagram on the right (Down DEGs by Both) shows genes downregulated in Both relative to anti-CD40 alone (Group D), anti-IgM alone (Group F), or both comparisons (Group E). Numbers indicate the gene count in each group. **C.** Gene expression trajectories for each group defined in Fig. 2B. **D.** Gene Ontology analysis for the ‘costimulation vs anti-CD40’ (Both vs CD40) and ‘costimulation vs anti-IgM’ (Both vs IgM) conditions at each stimulation time point. **E.** Gene expression trajectories representative of each group. Irf4 and Pdcd1lg2 belong to Group A; Myc and Tnfaip3 (A20) belong to Group B; Fas belongs to Group C; and Bcl6 belongs to Group D. **F.** Immunoblot of Blimp1 at the 24-hour time point. Splenic B cells from mice were stimulated as in Fig.1 over the indicated time period. The IAP inhibitor added 10 min after costimulation (+AT). Whole cell lysates were analyzed by immuno blotting with anti-Blimp1 Abs (n = 2).

We then asked whether cIAP shapes the Both-stimulated transcriptome, using the same 10-min post-stimulation inhibitor design validated in Fig. 1. The expression profiles of the ‘Both’ group and the ‘Both + IAP inhibitor’ group were classified into eight clusters based on their temporal trajectories (Fig. 3A, B). Clusters 4, 5, and 8 declined similarly under Both and Inh conditions and did not yield significant Gene Ontology enrichment, consistent with a largely IAP-independent, baseline-associated expression pattern; we do not characterize them further here. IAP inhibition caused the disappearance of late-phase (180 min) inflammation/cytokine induction via the canonical NF-κB pathway (Cluster 1: Ccl3, Ccl4, Cxcl10, Il10) and attenuated the early-phase NF-κB/AP-1 module (Cluster 7: Tnf, Icam1, Fosl1) (Fig. 3C). This is concordant with the IKK/ERK-duration-shortening logic we previously proposed (*10*). It has been shown that cIAP1/cIAP2 act in concert with TRAF2 as positive regulators necessary for canonical NF-κB activation downstream of the TNF receptor superfamily (including CD40) (*23*). Both the induction of acute-phase leukocyte activation and adhesion molecules, which peaks 60 minutes after costimulation, as seen in cluster 7,and the sustained antiviral/IFN-β response gene sets that intensify over time, as observed in cluster 1, are highly likely to constitute this canonical NF-κB-dependent transcriptional program; it is thought that the induction in both clusters has almost entirely disappeared as a result of this pathway becoming dysfunctional due to IAP inhibitors (Fig. 3D). In particular, the sustained IFN-β response in cluster 1 may lie downstream of a self-amplifying type I interferon loop initiated by early NF-κB activation; given that this loop itself cannot be initiated without the upstream canonical NF-κB pathway functioning, this is consistent with a simple ‘flattening’ trajectory. Spry2 itself, a negative regulator of B-cell proliferation mediated by BCR that is epigenetically silenced in B-cell lymphoma (*24*),was one of the four feedback regulators we initially identified (*10*), and it was included in this IAP-dependent group. Unexpectedly, the entire BCR-dominant module identified in Fig. 2 was also strongly cIAP-dependent (Fig. 3C): Cd5, Il10, Nfkbid (IκBNS, required for TACI expression and normal plasma cell differentiation; (*25, 26*)), and Spry1/2, none of which required CD40 costimulation. This provides a direct, data-driven candidate answer to the question we previously left open regarding which molecules mediate IAP’s action downstream of BCR. It also argues that cIAP is not a CD40-restricted adaptor but a hub shared by both receptors. In contrast, another set of genes (Cluster 2: Il6, Tnfsf4, Il12b, Trib3, Socs1) was gradually amplified by IAP inhibition. Their dynamics were consistent with, though not proof of, the release of NIK-dependent noncanonical NF-κB signaling. cIAP1/cIAP2 also act as ‘brakes’ that suppress the noncanonical NF-κB pathway (NIK–RelB/p52) under normal conditions, and it is known that IAP inhibitors lift this suppression to activate the noncanonical NF-κB pathway (*27–29*). Furthermore, experiments using cancer cell lines have reported that Smac mimetics elicit a biphasic response, in which a first wave of gene induction mediated by the canonical NF-κB/AP-1 pathway is followed by a second wave of gene induction mediated by TNFR1 signaling (*30*). The characteristic trajectory observed in cluster 3 in Fig. 3B (arrow head), ‘a group of genes peaking at the 30-minute mark appears, followed by a group peaking at the 60-minute mark’ corresponds very closely to this biphasic response (first wave + TNFR1-dependent second wave). Cluster 6, a smaller gene set that peaked transiently around 60 min under Both stimulation but, unlike the Both condition, failed to resolve toward baseline under IAP inhibition, shows a similar impaired-resolution kinetic to clusters 2 and 3; its gene set did not reach significant Gene Ontology enrichment (Fig. 3D), and we therefore treat it as supportive of, rather than separately diagnostic for, the same IAP-dependent de-repression pattern. The trajectory observed in cluster 2, ‘does not respond on its own but is monotonically and strongly induced in the presence of Inh’, can also be interpreted as consistent with the notion that the noncanonical NF-κB pathway, which is normally suppressed, has been activated as a result of IAP inhibition. This pathway has been separately suggested to be involved in the synergistic induction of Aicda/AID by BCR–TLR (*31*), which falls outside the scope of the original IKK/ERK duration model. We regard this as a hypothesis-generating finding, because our dataset did not detect target genes of the canonical or noncanonical pathways (Cxcl12, Cxcl13, Ccl19, Ccl21, Tnfsf13b), and because direct B-cell-specific support for individual genes in Cluster 2 is limited. Viewed together, the opposite consequences of IAP inhibition on Cluster 1/7 versus Cluster 2 delineate a dual, node-specific function for cIAP, sustaining canonical output while restraining non-canonical output at the same receptor-proximal complex, which we summarize as a working model in Fig. 3E.

**Fig. 3.**
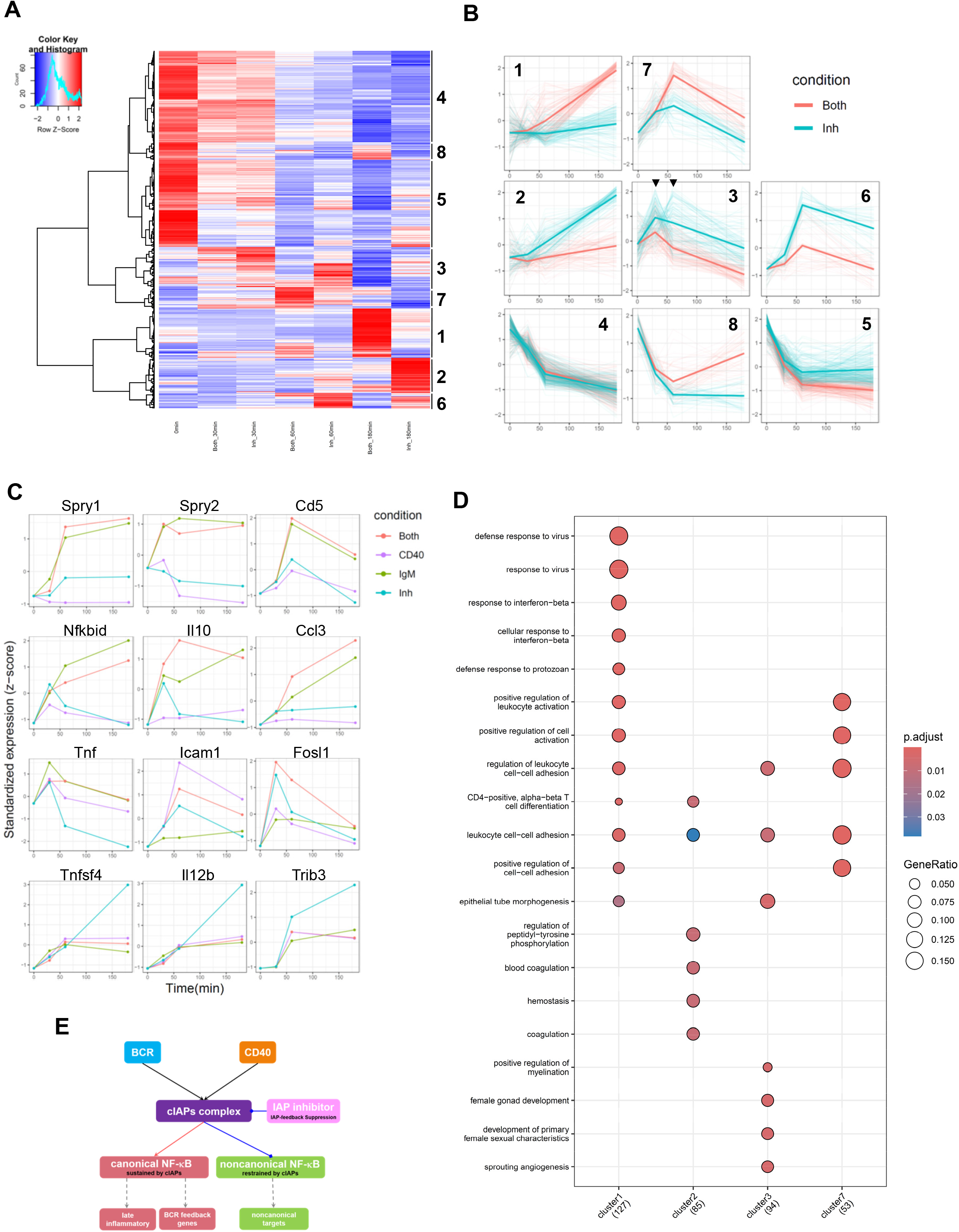
cIAP is required for the BCR-feedback module and restrains a delayed, non-canonical-NF-κB-like gene set. **A.** A heatmap of DEGs detected between B cells treated with anti-IgM and anti-CD40 (Both) and B cells treated with an IAP inhibitor 10 minutes after ‘Both’ stimuli (|log2FC| > 1.0). The cluster numbers are shown on the right-hand side of the heatmap. **B.** Gene expression trajectories for each cluster. The red line (Both) represents costimulation with anti-IgM and anti-CD40, whilst the blue line (Inh) represents the addition of an IAP inhibitor 10 minutes after costimulation. The numbers inside the boxes indicate the cluster numbers. **C.** Trajectories of gene expression belonging to each cluster. Spry1/2, Cd5, Nfkbid, Il10 and Ccl3 belong to Cluster 1; Tnf, Icam1 and Fosl1 belong to Cluster 7; and Tnfsf4, Il12b and Trib3 belong to Cluster 2. **D.** A Gene Ontology analysis for each cluster (1, 2, 3 and 7) defined in Fig. 3A. **E.** A proposed model in which BCR and CD40 converge on cIAP as a shared signaling hub: cIAP feedback sustains canonical, late-stage and BCR-specific negative-feedback gene expression while restraining output of the non-canonical NF-κ B pathway, thereby linking receptor-proximal signal dynamics to downstream cell-fate gene expression.

Together, these data extend our earlier demonstration (*10*) that IAP prolongs kinase activity after single-pathway stimulation, confirmed here under costimulation (Fig. 1). They further show that cIAP integrates BCR and CD40 inputs at the transcriptional level, supporting both the canonical late-gene program and, unexpectedly, the BCR-intrinsic negative-feedback module, while apparently restraining a non-canonical-NF-κB-like gene set. It is important to note that this study aims to identify correlations. Because both the RNA-seq data and the 24-hour Blimp1 data represent population averages, no conclusions can be drawn regarding fate differentiation at the single-cell level. Furthermore, while we previously classified Spry2 as part of the CD40-side feedback module (*10*), the present costimulation data classify Spry1/2 as belonging to the BCR-dominant class. One possible reason for this discrepancy is interspecies differences between mice and chickens. By inhibiting cIAP feedback, this study reveals, in part, the emergent nature of BCR and CD40 signaling. This finding should be useful for evaluating the efficacy of IAP inhibition for disease treatment. More broadly, these results indicate that a single feedback node, rather than two pathway-specific regulators, sets the duration of BCR and CD40 signaling and, through that duration, the transcriptional and fate output of the B cell, the defining property of a shared signaling hub.

## Methods

### Mice, cells, antibodies (Abs), and reagents

C57BL/6 mice from Charles River Laboratories International, Inc. were maintained under specific pathogen-free conditions and used at 8–12 week of age. All protocols were approved by the RIKEN Animal Committee and all experiments were carried out in accordance with the approved guidelines. For immuno blots, splenic B cells were purified by depleting CD43^+^ cells with magnetic beads using AutoMACS (Miltenyi Biotec) and were cultured in Iscove’s modified Dulbecco’s medium supplemented with 10% fetal bovine serum and 1% penicillin-streptomycin.

Abs specific for ERK, anti-phospho-ERK and anti-phospho-IκBβ were purchased from Cell Signaling Technology. Anti-PRDM1/BLIMP1 Antibody (EB05710) was from Everest Biotech. Anti-mouse IgM mAb (α-μ) was obtained from Jackson Immuno Research, and anti-mouse CD40 from BD Biosciences. The IAP inhibitor, AT, (AT-406; Active Biochem) was used at concentration of 10 μM.

For protein detection, the ECL Plex fluorescent western blotting system and ImageQuant LAS 4000 (GE Healthcare) were used. Kinase activity was quantified from the intensities of the protein and phosphorylated protein bands using a Multi Gauge version2.2 (Fujifilm) densitometer as described previously(*10, 32*).

### RNA sequencing, data preprocessing, alignment, and quantification

Total RNA from murine splenic B cells purified by depleting CD43+ cells was collected by using the NucleoSpin RNA kit (MACHEREY-NAGEL GmbH & Co.) according to the manufacturer’s instructions. For gene expression analysis, an Illumina library was prepared using a TruSeq Stranded mRNA and Truseq SR Cluster Kit v3 (Illumina, San Diego, CA) according to kit instructions. Sequencing was performed on a HiSeq 2000 sequencer (Illumina).

Raw 51-bp single-end RNA-seq reads were quality-filtered using PRINSEQ (version 0.20). The filtered reads were aligned to the Mus musculus reference genome (UCSC mm10 assembly) using TopHat2 (version 2.0.5). Gene-level read counts were subsequently generated from the aligned BAM files using featureCounts, implemented in the Subread package (version v1.3.0), with the corresponding gene annotation file (NCBI mouse RefSeq annotation (build 37.2)). Reads were summarized at the gene level based on the gene_id attribute. The resulting raw count matrix was used for subsequent bioinformatics analyses.

### Differential expression analysis

Differential expression analyses were performed in R (version 4.4.0) using the edgeR package (version 4.4.2). For comparisons between each stimulation time point (30, 60, and 180 min) and the unstimulated condition (0 min), genes with a maximum RPKM value greater than 1 in the relevant comparison were retained. Lowly expressed genes were further removed using the filterByExpr function. Count data were normalized for differences in library size using the trimmed mean of M-values method implemented in calcNormFactors. Dispersion parameters were estimated using estimateDisp and estimateGLMRobustDisp. Quasi-likelihood negative binomial generalized linear models were fitted using glmQLFit, and differential expression was assessed using the quasi-likelihood F-test implemented in glmQLFTest. The Benjamini–Hochberg method was used to control the false discovery rate (FDR), and genes with an FDR below 0.01 were considered differentially expressed.

For comparisons of the combined-stimulation condition with the IgM-stimulated, CD40-stimulated, or IAP inhibitor-treated condition at the corresponding time points, analyses were restricted to the union of genes identified as differentially expressed relative to the unstimulated condition. Genes exhibiting an absolute log2 fold change greater than 1 were selected as genes showing condition-dependent expression changes.

### Gene Ontology enrichment analysis

Gene Ontology (GO) enrichment analysis was performed using the clusterProfiler package (version 4.14.6) and the mouse annotation database org.Mm.eg.db (version 3.20.0). Gene symbols were used as identifiers. Upregulated and downregulated genes were analyzed separately at 30, 60, and 180 min using the compareCluster function with enrichGO. Enrichment analysis was restricted to the Biological Process ontology. P values were adjusted for multiple testing using the Benjamini–Hochberg method, with a nominal P-value cutoff of 0.05 and a q-value cutoff of 0.20. GO enrichment was also performed for selected gene clusters obtained from the heatmap analysis. Enrichment results were visualized as dot plots using clusterProfiler.

### Heatmap and hierarchical clustering

Heatmaps were generated from mean RPKM values using the heatmap.2 function in the gplots package (version 3.3.0). Expression values were standardized separately for each gene across the included conditions and time points by row-wise scaling. Genes were hierarchically clustered using correlation-based distance calculated with the amap package (version 0.8.20) and Ward’s minimum variance method (ward.D2). Column clustering was disabled, and the resulting gene dendrogram was displayed using a blue-to-red color scale. For the comparison between the combined-stimulation and inhibitor-treated conditions, the gene dendrogram was divided into eight expression clusters using the cutree function.

### Time-course visualization

Time-course expression profiles were visualized at 0, 30, 60, and 180 min using ggplot2 (version 4.0.3). RPKM values were standardized within each gene across the included conditions and time points. For gene sets and heatmap-derived clusters, the standardized expression profile of each gene was plotted as a semi-transparent line, and the mean standardized expression at each time point was superimposed as a thicker line. Expression profiles of selected individual genes were visualized using line and point plots. The number of differentially expressed genes at 30, 60, and 180 min was also summarized and displayed as a time-course line plot.

## Acknowledgments

We thank Dr. Shigehiro Kuraku for RNA sequencing in Phyloinformatics Unit, RIKEN Center for Life Science Technologies. We thank Anthropic’s Claude Sonnet 5 for assistance with manuscript preparation, including language editing, proofreading, and cross-checking of internal consistency between the text and figures. The authors take full responsibility for the content of this publication.

## Author contributions

H.S. designed and performed the experiments, K.I. performed the informatics analysis of transcriptome data, H.S., and K.I. wrote the manuscript.

